# Re-Targeting of Macroh2A Following Mitosis to Cytogenetic-Scale Heterochromatic Domains

**DOI:** 10.1101/333468

**Authors:** Hanae Sato, Bin Wu, Fabien Delahaye, Robert H. Singer, John M. Greally

## Abstract

The heritability of chromatin states through cell division is a potential contributor to the epigenetic maintenance of cellular memory of prior states. The macroH2A histone variant has properties of a regulator of epigenetic cell memory, including roles controlling gene silencing and cell differentiation. Its mechanisms of regional genomic targeting and maintenance through cell division are unknown. Here we combined *in vivo* imaging with biochemical and genomic approaches to show that human macroH2A is incorporated into chromatin in the G1 phase of the cell cycle following DNA replication. The newly-incorporated macroH2A re-targets the same, large heterochromatic domains where macroH2A was already enriched in the previous cell cycle. It remains heterotypic, targeting individual nucleosomes that do not already contain a macroH2A molecule. The pattern observed resembles that of new deposition of centromeric histone variants during the cell cycle, indicating mechanistic similarities for macrodomain-scale regulation of epigenetic properties of the cell.

## HIGHLIGHTS

- Unlike canonical histones, macroH2A is incorporated after DNA replication during G1.
- MacroH2A is organized into in large domains of hundreds of kilobasepairs.
- New macroH2A re-targets the loci of pre-existing macroH2A after DNA replication.
- MacroH2A is enriched in heterochromatic cytogenetic G-bands in the human genome.
- We find no evidence for homotypic organization of macroH2A in human nucleosomes.
- Approximately one human nucleosome in every eight contains a macroH2A molecule.

## INTRODUCTION

Epigenetic properties of cells are those involving differentiation decisions and memories of past events (Lappalainen and Greally, 2017). These properties are believed to be mediated at the molecular level by a number of transcriptional regulatory mechanisms. A necessary property of these epigenetic regulators of transcription is that they remain targeted to the same genomic regions in daughter chromatids following cell division, and only change with cellular differentiation. The replication of DNA introduces unmodified nucleotides, creating daughter chromatids with hemi-methylation of cytosine, the presence of 5-methylcytosine (5mC) on the template strand but not the complementary, newly-synthesized strand. This transient, hemi-methylated state is recognized and targeted for enzymatic re-establishment of 5mC on both strands (Bostick et al., 2007; Sharif et al., 2007). DNA replication also disrupts the association of proteins with DNA as the replication fork passes through a region, using pre-existing histones as well as freshly-synthesized histones that lack the post-translational modifications (PTMs) of the parent nucleosome to form new nucleosomes (Xu et al., 2010). While DNA methylation has a well-described biochemical mechanism for heritability through cell division, it has been more difficult to demonstrate comparable mechanisms for self-propagating maintenance of chromatin states. The existence of such mechanisms is supported by observations of self-propagation of the histone H3 PTM lysine 27 trimethylation (H3K27me3) in daughter cells over multiple cell divisions, despite the inactivation of the polycomb repressive complex 2 (PRC2) which catalyzes this PTM, in *Caenorhabditis elegans* (Gaydos et al., 2014) and in *Drosophila melanogaster* (Coleman and Struhl, 2017). Comparable findings have been revealed using nascent chromatin capture (NCC) and amino acid isotope labeling experiments (Alabert et al., 2015). The targeting of H3K27me3 in *D. melanogaster* appears to require the presence of polycomb-response elements (PREs) (Laprell et al., 2017), which mediate sequence-specific targeting by binding transcription factors (TFs) which then recruit the PRC2 complex. As we have previously noted (Henikoff and Greally, 2016), a model for the self-propagation of H3K27me3 is based on the ability of PRC2 to bind specifically to this modification (Hansen et al., 2008), suggesting that this binding tethers the PRC2 complex so that it can then add H3K27me3 onto other nearby nucleosomes post-replication.

There are other reasons why chromatin states can be the same in parent and daughter cells, for example those that are likely to be targeted as secondary consequences of other genomic processes. The passage of RNA polymerase through a region while transcribing a gene is associated with the local enrichment of PTMs such as H3K36me3, mediated by direct interaction of the Set2 lysine methyltransferase with RNA polymerase (Kizer et al., 2005). Histone PTMs at short regulatory elements flanking nucleosome-free regions are plausibly mediated by the recruitment of enzymatic complexes by transcription factors (Henikoff and Greally, 2016), while short RNAs such as the piwi-interacting RNAs (piRNAs) have been found to direct local repressive chromatin states at transposable elements in *D. melanogaster* (Le Thomas et al., 2013). More difficult to understand mechanistically has been the formation and maintenance through cell division of large chromatin domains exceeding tens of kilobases. Domains of this magnitude include the mediators of certain long-term cellular memories, such as the inactivation of an X chromosome during dosage compensation at the blastocyst stage of mammalian development (Augui et al., 2011), or the imprinting of large genomic domains during gametogenesis (Ferguson-Smith, 2011). Some of these larger-scale chromatin states involve the deposition of histone variants into nucleosomes in those regions. Histone variant deposition can be very focal, such as histone H3.3, which is enriched at *cis*-regulatory sites and telomeres (Goldberg et al., 2010), but others are maintained in broad genomic regions, such as CENPA at centromeric chromatin, occupying regions up to several million contiguous basepairs in size (Cleveland et al., 2003), and propagating to daughter chromatids through processes that are increasingly well understood (reviewed in (Müller and Almouzni, 2014)). The histone variant macroH2A also forms broad chromatin domains of at least hundreds of kilobases (Gamble et al., 2010) but is not limited to a discrete chromosomal location like the centromere, instead distributing genome-wide. MacroH2A differs from canonical H2A by having an additional C terminal ∼25 kDa globular domain (Pehrson and Fried, 1992), and has been shown to have roles both in the maintenance of cell states and in cell fate decisions (Barrero et al., 2013; Creppe et al., 2012; Pasque et al., 2012). The presence of macroH2A locally in the genome is mostly associated with transcriptional silencing, with striking enrichment at the inactive X chromosome territory in mammalian cells (Chadwick and Willard, 2002; Costanzi and Pehrson, 1998). Unlike other histone variants (Gurard-Levin et al., 2014), chaperones that target macroH2A to chromatin have not yet been identified, except in the specific situation of DNA damage, which involves macroH2A interacting with the Aprataxin-PNK-like factor (APLF) (Mehrotra et al., 2011). The loss of ATRX in the cell has been associated with a more permissive distribution of macroH2A into the alpha globin domain (Ratnakumar et al., 2012), suggesting that ATRX normally prevents the association of macroH2A with chromatin in this genomic region. Therefore, while we can implicate macroH2A as a potential contributor to the epigenetic property of cellular memory involved in X inactivation, we lack insight into how macroH2A propagates its genomic organization faithfully from parent to daughter cells, targeting specific regions of the genome, thus prompting the current study.

## RESULTS

To gain insights into how macroH2A remains targeted to specific genomic contexts through mitotic cell division, we performed a combination of imaging, biochemical and genomic techniques. We established the SNAP labelling system (Gautier et al., 2008) for macroH2A1.2 in HEK293T cells. As HEK293T cells have three X chromosomes, of which two are inactivated, the labelled macroH2A generates two strong X chromosome territory signals in each nucleus, a valuable marker of the stability and homogeneity of the modified cell line (**Figure S1** and supporting online **Video 1**). We used SNAP labelling combined with cell synchronization to demonstrate that we could distinguish prior macroH2A from newly incorporated macroH2A in a subsequent cell cycle, using separate fluorophores (**Figure S2A**). HEK 293T cells in prometaphase were collected following 12 hours of nocodazole blocking by mitotic shake-off, labelling the macroH2A present from the preceding cell cycle with SNAP-Oregon Green. The cells were then released from arrest at the G2/M transition, pulsed with SNAP-block to prevent any unconjugated macroH2A from detection by fluorophores, and returned to cell culture, arresting the cells at the next G2/M transition using RO-3306. Newly-incorporated macroH2A was detected with a distinctive red fluorophore using SNAP-TMR Star. This allowed live cell imaging during mitosis to be performed, measuring the levels of Oregon Green and TMR-Star in the dividing cells every 15 minutes (**Figure S2B**). The result showed that pre-existing macroH2A remained associated with chromatin following mitosis, and that both pre-existing and newly-incorporated macroH2A were evenly distributed into each of the daughter cells (**Figure S2C**).

We then tested the timing of incorporation of macroH2A. First, we determined incorporation timing of endogenous macroH2A1 through cell cycle by Western blot. Cells were harvested before and after S phase, a Western blot showing the expected increases in replication-dependent incorporation of histone H3 (Xu et al., 2010) and mitosis-associated enrichment of histone H3 serine 10 phosphorylation (Van Hooser et al., 1998) but little change in macroH2A1 (**Figure 1A**). We then applied an imaging-based approach to address the same question, using the cell synchronization and SNAP labelling approach shown in **Figure 1B**. We also added labelling with 5-ethynyl-2-deoxyuridine (EdU) as a further means of confirmation that cells were in the S phase of the cell cycle. We quantified the ratio of red (new) to green (old) histone in each cell for SNAP-H3 and SNAP-macroH2A (examples shown in **Figure 1C**). We demonstrated that the cells incorporated significantly more SNAP-H3 during S/G2 compared with the G1 phase (**Figure 1D**), once again consistent with the property of histone H3 being incorporated into chromatin during replication (Xu et al., 2010). By contrast, SNAP-macroH2A showed a distinctive pattern of strong enrichment in G1 but not S/G2 (**Figure 1D**), indicating that its incorporation was not dependent upon ongoing DNA replication.

**Figure 1:**
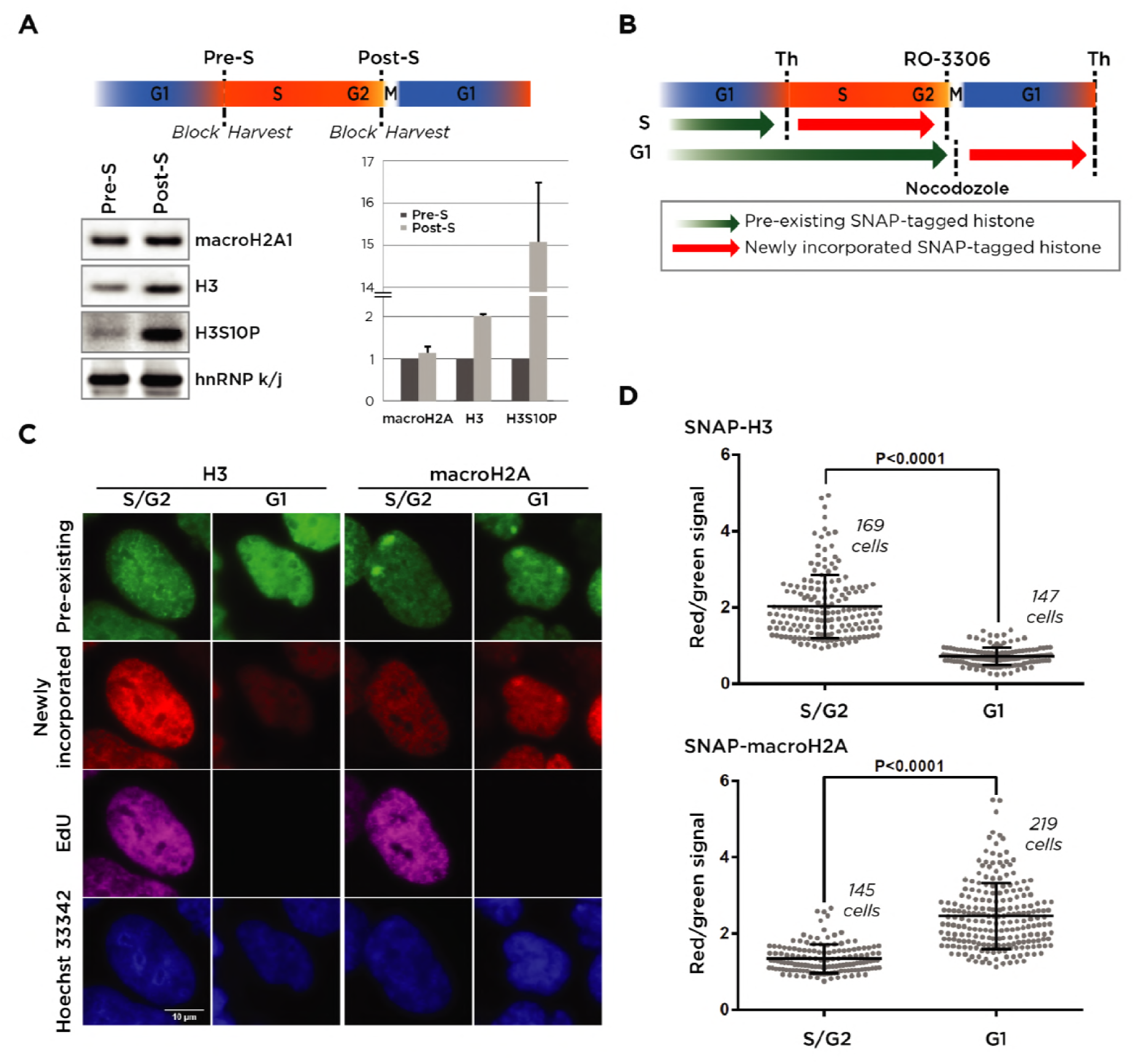
Quantification of SNAP-macroH2A in S/G2 and G1 phases of the cell cycle. **(A)** The time line of synchronization and harvesting of HEK 293T cells is shown. Equal numbers of cells were synchronized at the G1/S phase border by a double thymidine block. The cells were harvested (Pre-S) or released from thymidine block and synchronized prior to M phase with RO-3306 and harvested (Post-S). Chromatin fractions were isolated for Western blotting, testing endogenous macroH2A1, histone H3, and phosphorylation of H3 at serine 10 (H3S10P) as a marker of mitosis, with hnRNP k/j as a loading control. We show that H3 increase during mitosis, as expected, but there is no concurrent increase in macroH2A. n=3, Error bars = Standard deviation. **(B)** The time line of synchronization and labelling of cells at S/G2 and G1 phases for the analysis in (C). To label newly incorporated histones in S/G2 phase, HEK 293T cells stably expressing SNAP-tagged H3 or macroH2A were synchronized at the G1/S phase border by double thymidine block. Cells were treated with SNAP-Oregon Green to label pre-existing histones (green arrow), subsequently blocking non-labelled proteins using the non-fluorescent SNAP-Block reagent. The cells were allowed to progress to the G2/M transition until they were blocked using RO-3306 (a CDK1/cyclin B1 and CDK1/cyclin A inhibitor), labelling newly-incorporated SNAP-tagged histones with SNAP-TMR Star (red arrow). To label newly incorporated histones in G1 phase, mitotic cells were collected by shake-off following nocodazole treatment for 12 hours and spread onto coverslips. After two hours, cells were labelled with Oregon Green and treated with the blocking reagent. Cells were then allowed to progress to the G1/S transition, when they were synchronized by double thymidine block, then labelling newly-incorporated SNAP-tagged histones with SNAP-TMR Star (red arrow). After being released from the first synchronization, the cells were also incubated with EdU until the second synchronization, allowing cells that had undergone DNA synthesis to be identified. **(C)** An example of images showing the detection of pre-existing and newly-synthesized SNAP-tagged histones in S/G2 or G1 phases. Bar = 10 µm. **(D)** Image analysis measurements of red and green nuclear signals, representing the ratio of newly-incorporated to pre-existing histones H3 and macroH2A in the S/G2 and in G1 phases. The error bars represent one standard deviation. The P values were determined using two-tailed unpaired t-tests.

While X chromosome inactivation can occur in the absence of macroH2A (Tanasijevic and Rasmussen, 2011), macroH2A1 appears to work synergistically with the PRC1 polycomb complex and the CULLIN3/SPOP ubiquitin E3 ligase to stabilize inactivation (Hernández-Muñoz et al., 2005), and appears to be recruited by the *Xist* long non-coding RNA (Csankovszki et al., 1999). The inactive X is also notable for its late replication timing during S/G2 (Koren and McCarroll, 2014), raising the question whether there is a distinctive pattern of deposition of macroH2A during the cell cycle in the inactive X chromosome compared with the rest of the genome. We partitioned the nuclear signal into the subnuclear domains containing the two inactive X chromosomes and the remainder of the nucleus. The inactive X territories were apparent from the pre-existing macroH2A signals. The relative signal from SNAP-macroH2A ithin these territories was compared with the remainder of the nucleus. We found the incorporation of macroH2A into the inactive X chromosomes occurred at the same time as the remainder of the genome, during the G1 phase of the cell cycle (**Figure S3**).

We then performed a time course experiment to gain more precise insights into the timing of deposition of macroH2A during the cell cycle. Cells were synchronized at the G1/S transition using a double thymidine block, and then released and cultured for up to 22 hours. Four fluorophores were used in these cells, two for SNAP labelling of macroH2A and two for the Fucci cell cycle sensor system (Sakaue-Sawano et al., 2008). The cells were then sampled every 2 hours (**Figure 2A**) and tested first using flow cytometry to quantify the intensities of signals of pre-existing SNAP-macroH2A (green) and newly incorporated macroH2A (red). We show in **Figure 2B** that the intensity per cell of pre-existing SNAP-macroH2A drops suddenly at 14 hours. Parallel studies using the Fucci cell cycle sensor system shows the HEK 293T cells to be in S/G2 until 12 hours with a change to G1 at 14 hours (**Figure S4B**). The decrease of signal intensity of pre-existing macroH2A therefore coincides with cells undergoing mitotic division and the dilution of the pre-existing macroH2A into two daughter cells. The signal for newly-incorporated SNAP-macroH2A, on the other hand, began to be observed at 18 hours, during the G1 phase.

**Figure 2:**
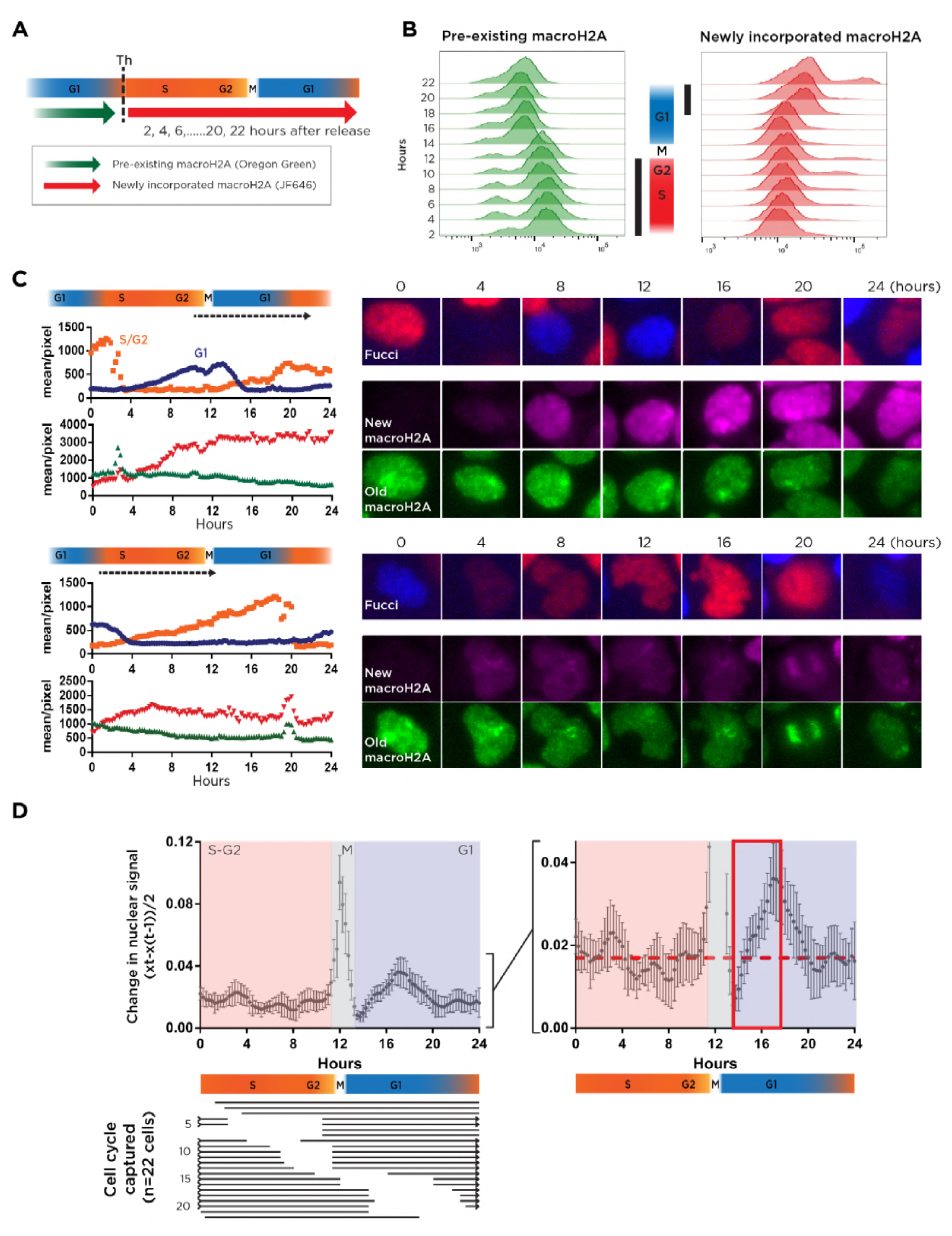
Flow cytometric and real time *in vivo* imaging of SNAP-macroH2A incorporation timing. **(A)** The time line of synchronizing and labelling cells. Newly synthesized macroH2A was labelled with SNAP substrate JF646 at each 2 hour time point after release from thymidine block. **(B)** Flow cytometric analysis of pre-existing (Oregon Green) and newly incorporated macroH2A (JF646) and the Fucci cell cycle indicator. The stage of the cell cycle at each time point was determined by the levels of TagBFP (G1 marker) and mCherry (S/G2 marker) (**Figure S4**), and indicated in the middle section. The black columns represent when pre-existing macroH2A (hours 2-12, S/G2) transitions to an enrichment for newly-incorporated macroH2A (hours 18-22, G1). **(C)** Examples of *in vivo* imaging of macroH2A dynamics in individual cells. Pre-existing macroH2A in non-synchronized cells was labelled with SNAP-Oregon Green and blocked with SNAP-Block. Newly-incorporated macroH2A was detected with JF646. Live cell images were acquired every 20 minutes for 18-20 hours. Signal intensity transitions were measured as shown in the left panels. Representative images from 4 hourly intervals are shown in the right. The upper cell is captured entering G1 (dashed arrow) and shows a substantial accumulation of new macroH2A, whereas the lower cell is captured during S/G2 and accumulates new macroH2A to a much lesser extent. Bar=10µm. **(D)** The summarized data from imaging of 22 cells. The cells were aligned temporally using the Fucci cell cycle images. The lines in lower panel represent the detection phase in each cell. The rate of change of signal of newly incorporated macroH2A was calculated as the difference of intensity of newly-incorporated macroH2A normalized to the signal in first time point of pre-existing macroH2A between two consecutive time points (Delta (X_t_-X_(t-_ 1))/2). These delta values were then plotted through the cell cycle. Signal saturation at metaphase distorts the data, but the right panel allows comparison of temporal changes compared with the average delta (0.17) during S-G2 phase. The period of sustained accumulation of macroH2A is between hours ∼13-17 (red box). Error bars = Standard error of the mean.

We complemented these flow cytometry studies with live cell imaging to gain more detailed resolution of the timing of acquisition of newly-incorporated SNAP-macroH2A. Representative results are shown in **Figure 2C** (and supporting online **Videos 2-3**), with the summary of the imaging of 22 cells in **Figure 2D**. We measured the signal intensity of newly-incorporated macroH2A, normalized by the signal from pre-existing macroH2A, and calibrated for each cell the stage of the cell cycle using the Fucci signals. We found that the single period of consistent macroH2A incorporation was between hours 13-17, starting immediately after metaphase and extending to mid-G1 phase (**Figure 2D**).

Knowing that new macroH2A was being incorporated during early G1, we could then ask whether it was being targeted at that time to the genomic regions where the macroH2A had been incorporated in the parent cell. We again exploited the SNAP labelling system to conjugate biotin to pre-existing and newly-incorporated macroH2A, as shown in **Figure 3A**. We performed the equivalent of native chromatin immunoprecipitation (N-ChIP) using SNAP-biotin instead of antibodies to enrich the subset of nucleosomes with SNAP-macroH2A. We were able to get enough enrichment from the S/G2 preparation from residual unblocked SNAP tags to represent the chromatin of the parent cell, with the expected higher yields from the G1 sample with its unblocked SNAP tags available for conjugation to biotin. The libraries were sequenced in parallel with an input sample.

**Figure 3:**
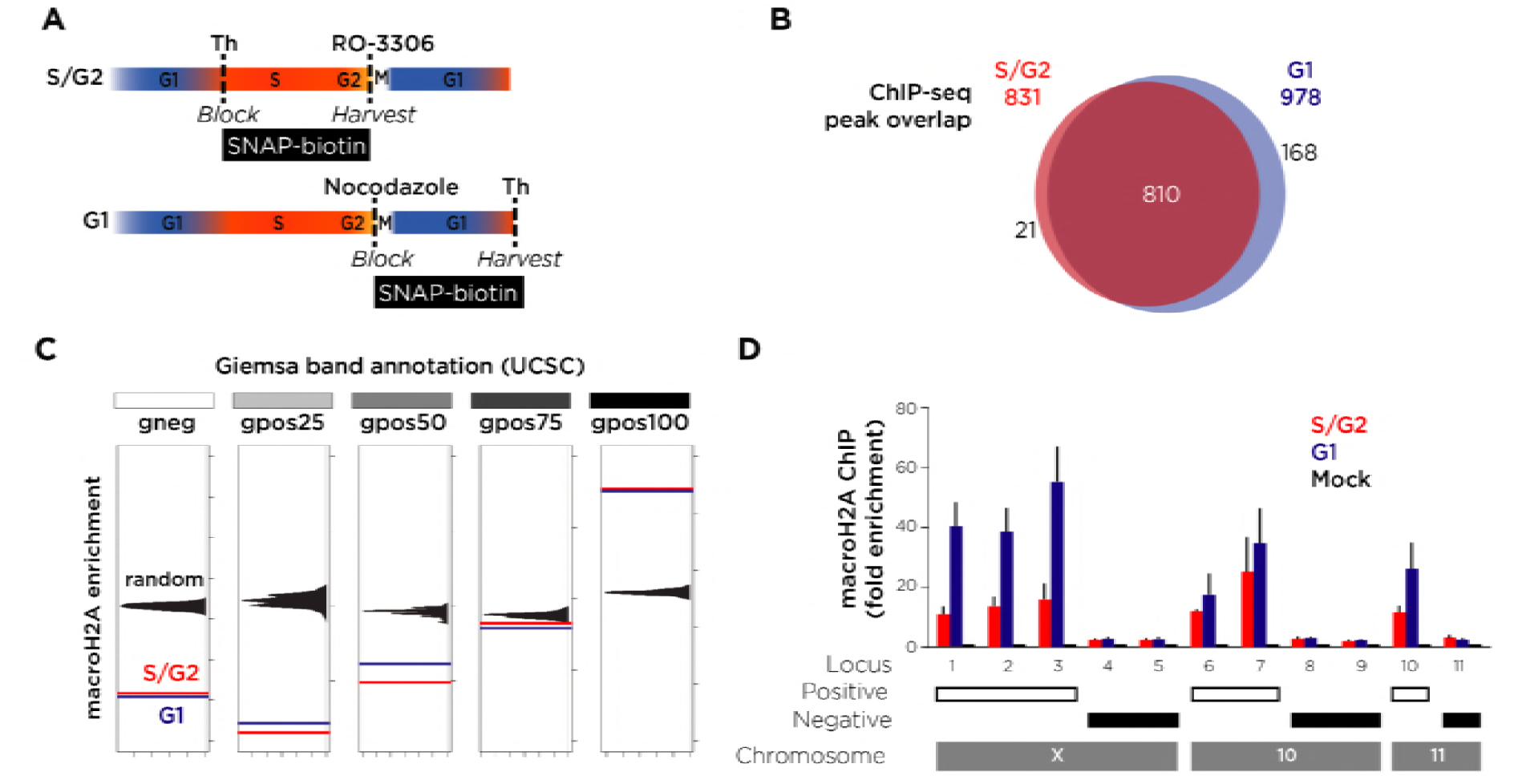
Cell cycle-specific ChIP-seq studies of macroH2A incorporation in S/G2 and G1 phases of the cell cycle. **(A)** The time line of synchronizing, blocking and harvesting of cells. **(B)** Using data from 500 kb genomic windows and identifying those with high confidence for macroH2A enrichment (**Figure S5**), we find that these “peaks” overlap substantially between macroH2A newly incorporated in S/G2 and in G1. **(C)** MacroH2A is enriched before and after cell division in Giemsa positive bands, specifically those categorized as gpos100 in the UCSC Genome Browser. The results of permutation tests demonstrate enrichment only for the most heterochromatic cytogenetic bands (P<0.01). **(D)** ChIP-qPCR of loci predicted from the ChIP-seq results to be positive and negative validates these genome-wide studies.

Previous studies mapping the locations of macroH2A in mammalian cells revealed it to be enriched in very broad domains, requiring modification of standard ChIP-seq peak calling approaches (Gamble et al., 2010; Yildirim et al., 2014). We therefore started our analysis by testing the enrichment patterns in genomic windows of different sizes (**Figure S5**). The expected ChIP-seq pattern should be of enrichment in some genomic locations relative to others, resulting in a bimodal distribution of sequencing reads from those loci when comparing affinity-purified against input mononucleosomal samples. Using windows of 1–1,000 kb, we observed the bimodal distribution indicating enrichment at ≥500 kb resolution. We defined the inflection point separating the loci of macroH2A enrichment for the S/G2 and G1 phases of the cell cycle using the *pastecs* R package (**Figure S6**). The enriched 500 kb windows identified in this way represented our highest confidence loci for macroH2A deposition in the genome, allowing us to test how concordant these loci were during G1 and S/G2 phases of cell cycle. Of the 831 windows with macroH2A enrichment during S/G2 phase, 810 (97.6% of windows in S/G2 phase) remained enriched in G1 phase at loci of newly-incorporated macroH2A (**Figure 3B**). Visual inspection of the results on a genome browser indicated that the windows of enrichment were located especially in Giemsa dark (G-) bands, which we confirmed through permutation studies of randomly redistributed windows of enrichment of macroH2A and UCSC Genome Browser annotations of cytogenetic bands (**Figure 3C**). Using quantitative PCR, we showed that the loci in windows with predicted macroH2A deposition were indeed enriched (**Figure 3D**). These genomic localization studies therefore showed macroH2A to be enriched in very large domains of hundreds of kilobases, especially in the cytogenetic G-bands representing constitutive heterochromatin, and that newly-incorporated macroH2A in G1 targets the loci already enriched for macroH2A in the parent cell.

The question that arose was how macroH2A recognizes these heterochromatic regions already enriched for this histone variant. In eukaryotic cells, the chromatin is organized by the basic unit of the nucleosome, which is composed by two dimers of H2A-H2B and a tetramer of H3-H4 in 147 base pairs of DNA. We tested the simplest possible model, that macroH2A recognizes individual nucleosomes that already contain both macroH2A and H2A heterotypically, and replaces the existing H2A with a second macroH2A molecule to create a homotypic nucleosome. We prepared mononucleosomes (**Figure 4A**) and used an anti-SNAP antibody that we had characterized (**Figure S7**) to isolate the subset of nucleosomes containing a SNAP-macroH2A (**Figure 4B**). Using an anti-macroH2A1 antibody, we tested whether these nucleosomes also included endogenous macroH2A, which would indicate the presence of nucleosomes homotypic for macroH2A. The HEK293T cell line used for all assays in this project (except where indicated otherwise) expressed SNAP-macroH2A at low levels (30% of endogenous macroH2A (**Figure S8A**)), enhancing the chances of finding homotypic nucleosomes containing endogenous macroH2A and SNAP-macroH2A. Endogeneous macroH2A could not be detected in the nucleosomes containing SNAP-macroH2A (**Figure 4B**).

**Figure 4:**
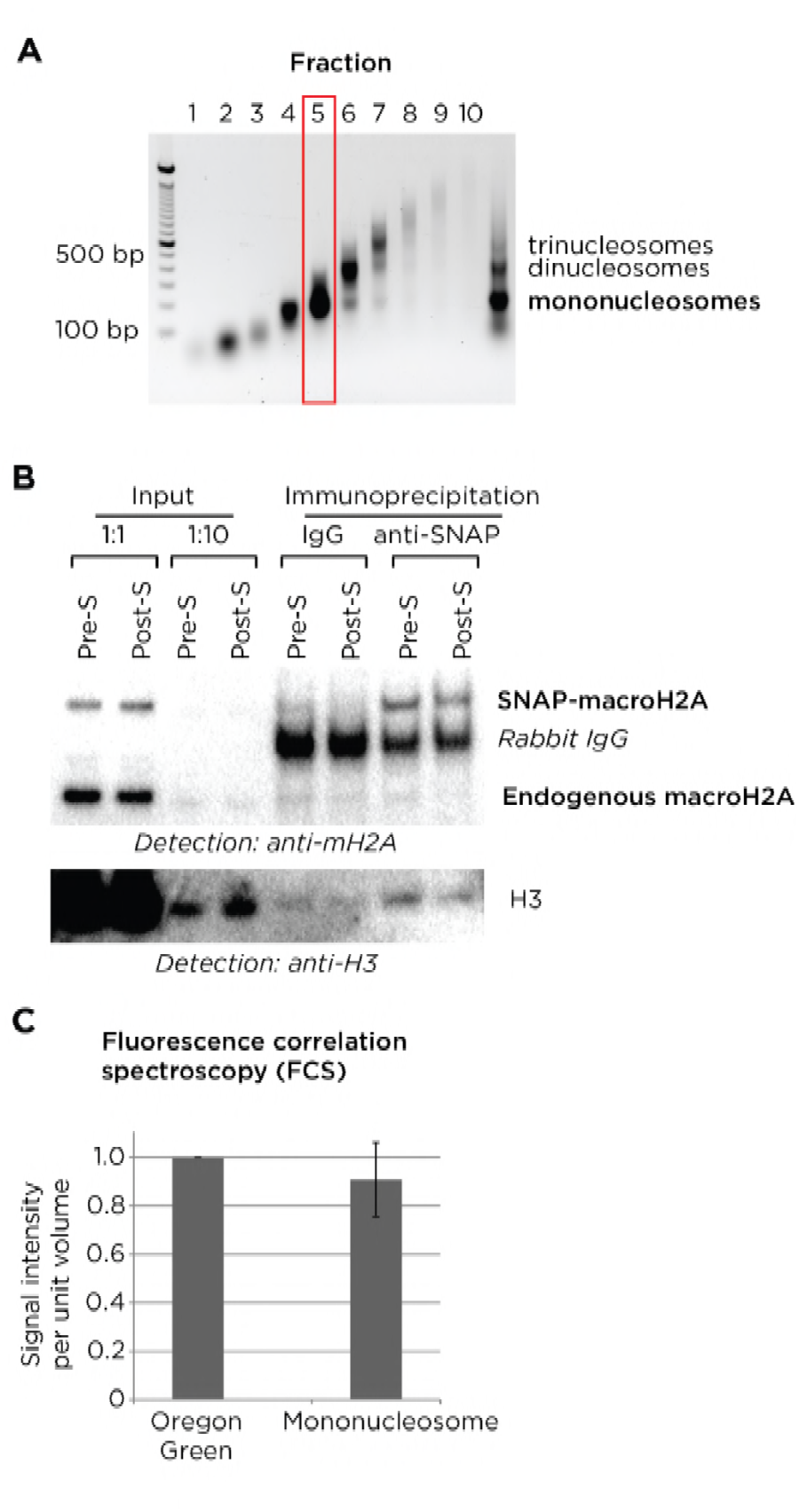
Nucleosomal organization of macroH2A. **(A)** Mononucleosomes (red box) were purified using high density sucrose gradient ultracentrifugation of samples synchronized as in Figure 1A. **(B)** Western blotting before and after immunoprecipitation of pure mononucleosomes using anti-SNAP antibody or rabbit IgG as a negative control. Input samples (and 1:10 dilutions) were loaded in the four lanes on the left. The lower Western blot shows that histone H3 is present in the isolated mononucleosomes and that the loading of the pairs of samples prior to or after S phase (indicated as Pre-S or Post-S) lanes are balanced. In the upper blot, there is no evidence for nucleosomes containing SNAP-macroH2A also containing detectable levels of endogenous macroH2A, as the signal intensities in the anti-SNAP lanes do not exceed those of the non-specific IgG lanes. **(C)** The brightness of individual nucleosomes containing Oregon Green-labelled SNAP-macroH2A measured by fluorescence correlation spectroscopy (FCS) was indistinguishable from individual beads with single molecules of SNAP-Oregon Green, demonstrating that individual nucleosomes contain only single molecules of SNAP-macroH2A.

We then applied fluorescence correlation spectroscopy (FCS) (Chen and Müller, 2007) to test this question in an orthogonal manner. FCS allows the quantification of fluorescence intensity of single molecules in solution, in this case allowing us to test whether the single nucleosomes isolated contained more than one SNAP-tagged molecule, indicating a homotypic organization of the histone variant. For these experiments we switched to a cell line expressing SNAP-macroH2A at a level three times higher than endogenous macroH2A1, increasing the chance of finding nucleosomes with two SNAP-macroH2A molecules in any homotypic nucleosomes present (**Figure S8A**). To ensure that we were saturating the labelling of SNAP-macroH2A, so that any subset of nucleosomes containing two SNAP-macroH2A molecules would reliably show two fluorescent molecules, we defined and used the saturating conditions for SNAP labelling (**Figure S8B**). We measured the fluorescence intensity per unit volume of a solution containing single nucleosomes saturated for SNAP-macroH2A labelling, compared with a solution containing individual SNAP-Oregon Green molecules. In **Figure 4C,** we show that the signal intensity per nucleosome is indistinguishable from that of single SNAP-Oregon Green molecules, demonstrating that only single SNAP-tagged macroH2A molecules are detected in individual nucleosomes. Our results are consistent with the prediction that macroH2A is likely to be unstable when present homotypically in a nucleosome (Chakravarthy and Luger, 2006), and exclude the possibility of incorporation of new macroH2A into nucleosomes already containing macroH2A as the mechanism of targeting macroH2A during G1 phase to genomic regions already enriched in this histone variant.

Finally, we estimated the proportion of nucleosomes in the human genome containing a macroH2A molecule. We used cell lines expressing either SNAP-macroH2A or SNAP-H3 and isolated mononucleosome preparations which were loaded onto a Western blot (**Figure S7**). Detection with an anti-H3 antibody allowed a loading control for mononucleosome numbers, showing the endogenous and SNAP-tagged H3 proteins. Detection with an anti-SNAP antibody revealed the difference in expression levels of the transgenes, while detection with anti-macroH2A showed us the relative expression levels of the transgenic and endogenous macroH2A genes. This in turn allowed us to estimate the proportion of nucleosomes in the SNAP-H3 cell line containing macroH2A, calculated as 12.1%. Genome-wide, it therefore appears that approximately one nucleosome in every eight contains a macroH2A histone variant, probably occurring at much higher proportions in G-bands, and lower proportions in the remaining majority of the genome.

## DISCUSSION

The combination of approaches used in this study have revealed insights into the heritability through cell division of a chromatin state that is organized over a scale of hundreds of kilobasepairs in the human genome. MacroH2A appears to occupy nucleosomes heterotypically with histone H2A, enriched in Giemsa dark G-bands, which are the most heterochromatic and late-replicating regions of the genome (Suzuki et al., 2011). These late-replicating regions in the genome are those targeted for deposition of new macroH2A in the hours following mitosis and cytokinesis, during the G1 phase in daughter cells. The mechanism for re-targeting of macroH2A to these regions is unknown, but does not involve recognition of and incorporation of macroH2A into individual nucleosomes already containing macroH2A. As H2A is already present in the nucleosomes formed during daughter chromatid formation in S/G2, the subsequent targeting of macroH2A involves replacement of one of the two H2A molecules in individual nucleosomes.

This timing of incorporation of the macroH2A histone variant into chromatin following cell division in the G1 phase reveals a parallel with the physiology of the centromeric histone variant CENP-A (Jansen et al., 2007). CENP-A is the H3-like histone variant that is part of the specialized nucleosome forming the centromere (French and Straight, 2013). Human centromeres are estimated to be up to several million contiguous basepairs in size (Clevel et al.), a magnitude of the same order as that inferred from our ChIP-seq data for macroH2A. There is a strong preference in human cells to form centromeres at specific short satellite DNA sequences, but they can also be formed ectopically at other sequences, and remain stable at those locations through cell division and across generations (Amor and Choo, 2002), representing a molecular mechanism for epigenetic maintenance of cellular memory. Following DNA synthesis and cell division, pre-existing CENP-A remains at centromeres but is distributed between daughter chromatids with H3.3 (Dunleavy et al., 2011). The subsequent re-targeting of these diluted locations with new CENP-A in G1 is of uncertain mechanism, but may involve histone H3 lysine 4 dimethylation (H3K4me2) and the activity of the HJURP chaperone for CENP-A (Bergmann et al., 2011), which is in turn recruited to the mammalian centromere by the Mis18 complex (Wang et al., 2014) assisted by a centromeric long non-coding RNA (Quénet and Dalal, 2014). These observations about the centromere offer potential guidance into how we might study the targeting of macroH2A.

Our observation that macroH2A is preferentially targeted to cytogenetic bands with the characteristics of heterochromatin is consistent with its known property to maintain heterochromatic structures. The association of macroH2A with the inactive X chromosome is well-known, but more recently it has also been demonstrated that when macroH2A was depleted in HepG2 cells, cytologic changes in heterochromatin and nucleolar organisation became apparent (Douet et al., 2017). The same study also found macroH2A to be associated with heterochromatic and H3K9me3-enriched regions of the genome, consistent with the findings presented here, and also demonstrated a role for macroH2A in the attachment of SAT2 repeats to Lamin B1 (Douet et al., 2017). These authors concluded that macroH2A plays a significant role in maintaining nuclear architecture, in particular the association of heterochromatic, H3K9me3-enriched regions with the nuclear lamina, shedding light on one aspect of its functional role in the cell nucleus. At the nucleosomal level, how macroH2A exerts its repressive effects appears to involve the direct interference by the macroH2A tail domain of NF-kappaB binding to DNA locally, and a resistance of chromatin containing macroH2A to SWI/SNF-mediated chromatin remodelling (Angelov et al., 2003). The repressive effects of macroH2A are therefore likely to be acting both at the level of individual nucleosomes and at the level of subnuclear organization of chromatin, implying that macroH2A is a multifunctional repressor of chromatin *in vivo*.

Despite the interest in macroH2A’s roles in cell state maintenance and cell fate decisions (Barrero et al., 2013; Creppe et al., 2012; Pasque et al., 2012), surprisingly little is known about how it is inherited through cell division. The landmark study on which most of our current insights are based was performed in 2002 using immunofluorescence techniques (Chadwick and Willard, 2002), mostly focused on the association of macroH2A with the inactive X chromosome. We note that our imaging results do not support their immunofluorescence-based observation of a dissipation of the macrochromatin body at the inactive X chromosome during late S phase and G2. On the other hand, their finding that macroH2A reforms following cell division during the G1 phase of the cell cycle is consistent with our imaging and biochemical results. The major value of the current study is to provide an updated fundamental set of observations about macroH2A heritability through cell division, a necessary foundation if we are to progress to the identification of chaperone-mediated mechanisms that help to target this histone variant to heterochromatin. The discovery of the targeting mechanisms for regional macroH2A deposition during G1 will represent a significant insight into the epigenetic mechanisms of cellular memory occurring on a macrodomain scale, a mechanistically under-explored area of research. We recognize the presence of comparably large domain organization in normal cells and in cells with regulatory perturbations, such as partially-methylated domains (Gaidatzis et al., 2014) and lamina-associated domains (Guelen et al., 2008), but we lack insight into how these large domains are co-ordinately regulated. The influences that maintain or perturb large-scale macroH2A targeting will provide valuable insights into transcriptional regulatory influences conferring epigenetic properties to the cell, acting over hundreds to thousands of kilobases.

## AUTHOR CONTRIBUTIONS

Conceptualization, H.S., J.M.G.; Methodology, H.S., B.W., R.H.S., J.M.G.; Formal Analysis, H.S., B.W., F.D.; Investigation, H.S.; Resources, R.H.S.; Data Curation, H.S.; Writing – Original Draft, H.S., J.M.G.; Writing – Review & Editing, H.S., R.H.S., J.M.G.; Visualization, H.S., J.M.G.; Supervision, R.H.S., J.M.G.; Project Administration J.M.G.; Funding Acquisition J.M.G.

## ACKNOWLEDGMENTS

This work was supported by NIH grant R01 DA030317 (JMG). We thank members of the Greally and Singer laboratories for discussions, Professor Noel Lowndes and Dr. Joseph McCarter (National University of Ireland, Galway) for advice on protocols, the Einstein Epigenomics Shared Facility and FACS and Genomics cores, and Einstein’s Center for Epigenomics. We also thank L. Lavis for SNAP-JF646 and Y. Kong for help with ChIP-seq analysis. The authors declare no conflicts of interest.

## VIDEOS

Uploaded separately.

**Video 1:** The live cell imaging of the images shown in **Figure S1B**.

**Video 2:** The live cell imaging of the images shown in **Figure 2C (top)**.

**Video 3:** The live cell imaging of the images shown in **Figure 2C (bottom)**.

## SUPPLEMENTARY MATERIALS

**This section includes:**

Supplementary Figures S1 to S9

Supplementary Table S1

Supplementary Methods

Supplementary References

## SUPPLEMENTARY FIGURES

**Supplementary Figure S1:**
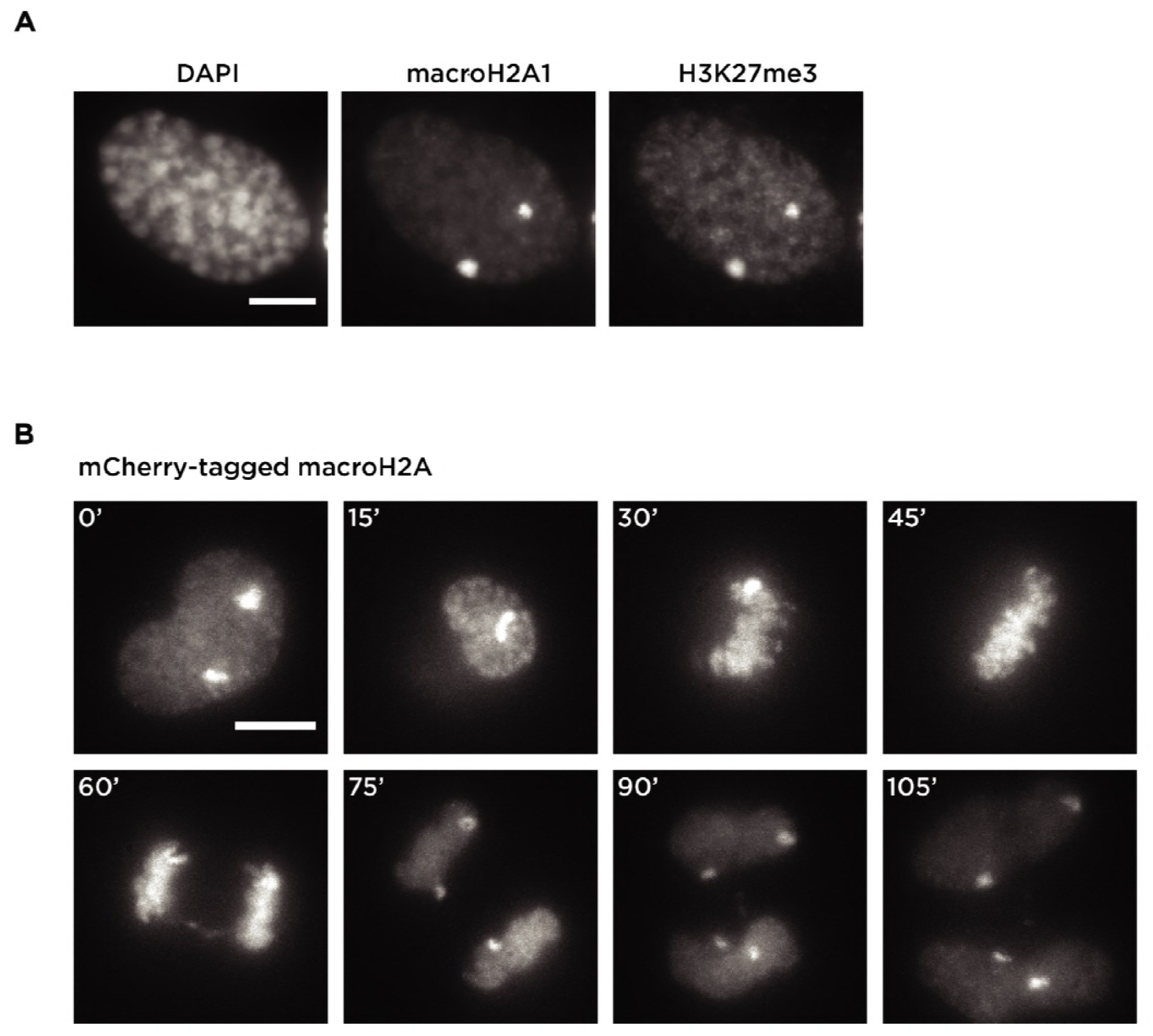
macroH2A remains associated with the inactive X chromosome during mitosis in HEK 293T cells. Immunofluorescence of endogenous macroH2A1 and histone H3 trimethylation (H3K27me3). Enrichment of endogenous macroH2A1 co-localized with H3K27me3 in the two inactive X chromosomes was detected at interphase in a HEK 293T cell. Bar=10 µm. **(B)** Live cell imaging of mCherry-tagged macroH2A during mitosis. mCherry-tagged macroH2A1.2 was expressed under a CMV promoter. A mitotic HEK 293T cell was imaged using widefield microscopy. Time series images were acquired every 15 minutes. HEK 293T cells contain three X chromosomes, with two inactivated, apparent as the two bright subnuclear signals of mCherry-tagged macroH2A. Bar=10 µm.

**Supplementary Figure S2:**
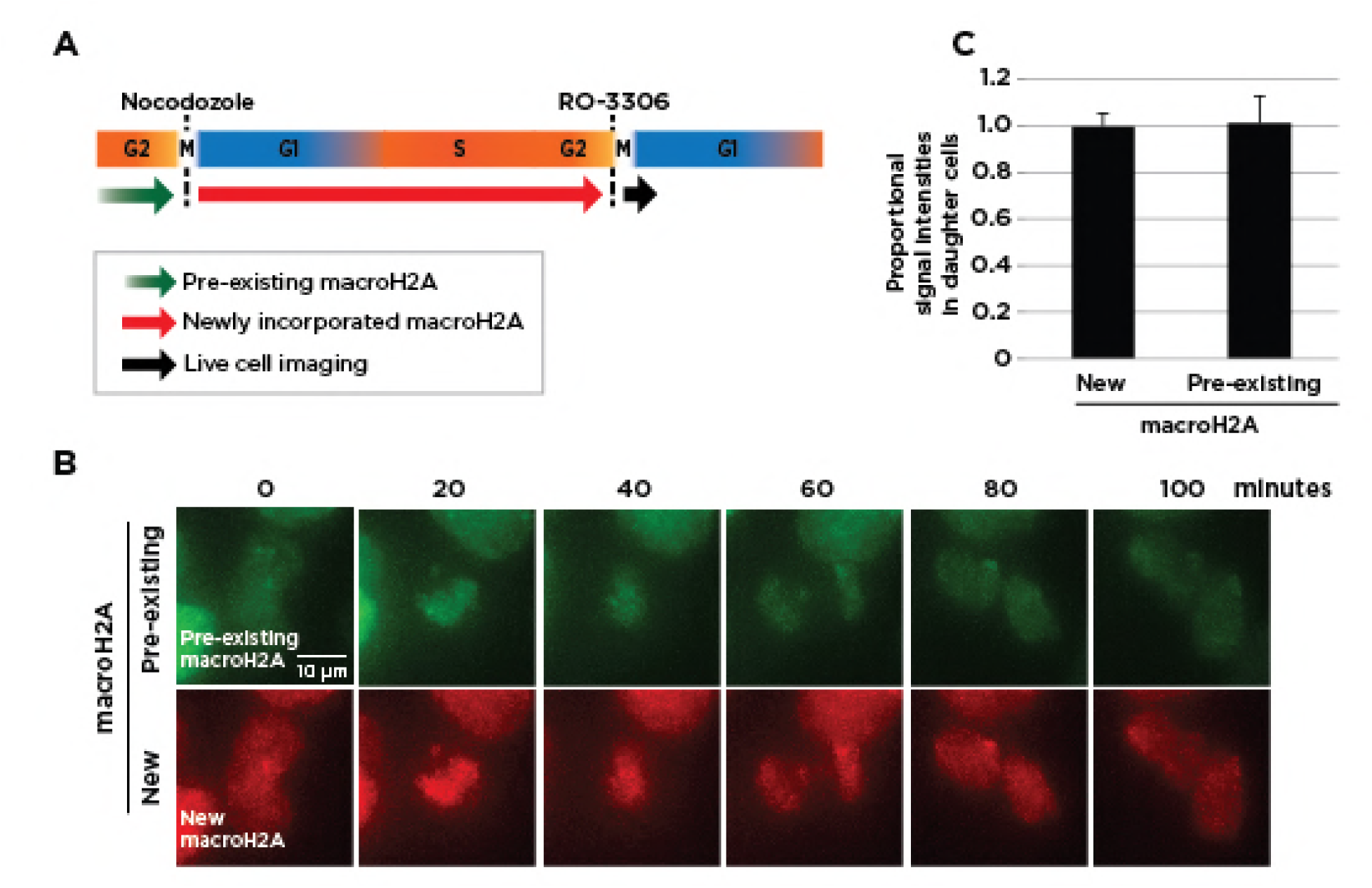
Live cell imaging of pre-existing and newly incorporated macroH2A distributions during mitosis. The timeline of synchronization and labeling of cells. Cells in prometaphase were collected by shake-off following nocodazole treatment for 12 hours. Three hours after spreading onto a MatTeck cell culture dish, the pre-existing SNAP-tagged macroH2A was labeled with SNAP-Oregon Green (OG) to cause the existing MacroH2A to fluoresce green. The cells were then pulsed with SNAP-Block to prevent any remaining SNAP-tagged macroH2A from being able to conjugate with fluorophores. Following further culture, the cells were arrested at the G2/M transition by RO-3306 treatment, allowing SNAP-macroH2A newly incorporated since the G2/M transition to be labeled with SNAP-TMR STAR (TMR) for a red fluorescence signal. Live cell imaging was then performed after releasing from the RO-3306 block. **(B)** Live cell imaging of pre-existing and newly incorporated SNAP-macroH2A distribution into daughter cells during mitosis. Pre-existing (OG, green) and newly incorporated (TMR, red) SNAP-macroH2A were imaged at 20 minute intervals. **(C)** *In vivo* imaging reveals equal distributions of pre-existing and newly incorporated SNAP-macroH2A to daughter cells (n=5).

**Supplementary Figure S3:**
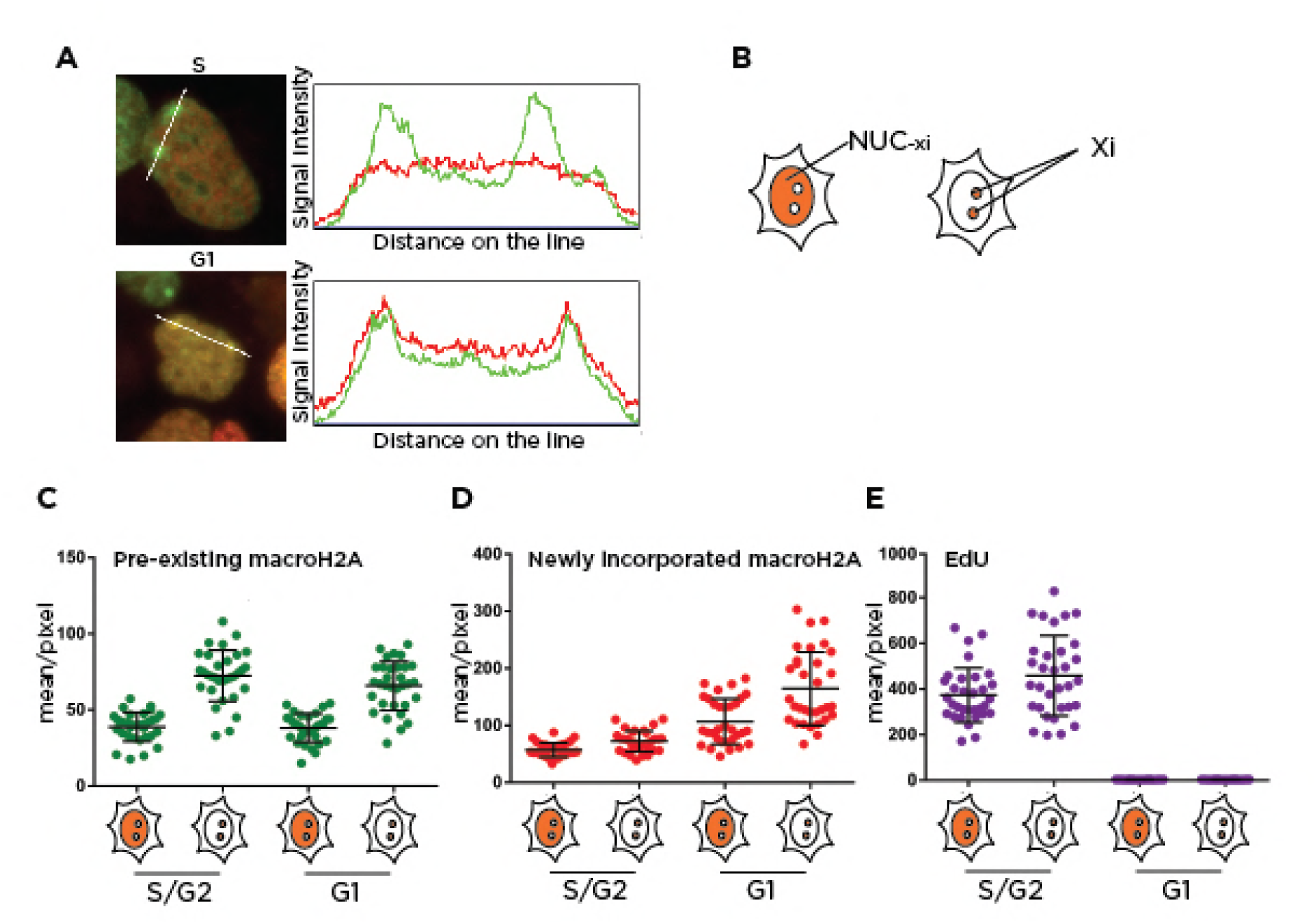
Pulse-chase detection of macroH2A incorporation on Xi. **(A)** Image analysis of SNAP-tagged macroH2A incorporation on the two inactive X chromosomes (Xi) in HEK 293T cells. A line of HEK 293T cells stably expressing SNAP-tagged macroH2A was synchronized and pulse-chase labeled as shown in **Figure 1B**. The left panels show the merged images of pre-existing (Oregon Green: green) and newly incorporated (TMR: red) SNAP-macroH2A. In the right panels we show the signal intensities measured along the white dashed lines in the left panels. The green and red lines show the signal intensities of pre-existing and newly incorporated macroH2A per pixel, respectively. **(B)** We show the areas measured. NUC_-Xi_ is the area of nucleus excluding the two inactive X chromosomes (Xi). **(C)** The signal intensities of pre-existing macroH2A in NUC_-xi_ and Xi. **(D)** Comparison of signal intensities of newly incorporated macroH2A between the NUC_-Xi_ and Xi subnuclear domains in S/G2 and in G1. **(E)** The signal intensities of EdU incorporation on NUC_-Xi_ and Xi. No differences are apparent between the dynamics of incorporation of newly-incorporated macroH2A into the inactive X chromosome compared with the rest of the nucleus.

**Supplementary Figure S4:**
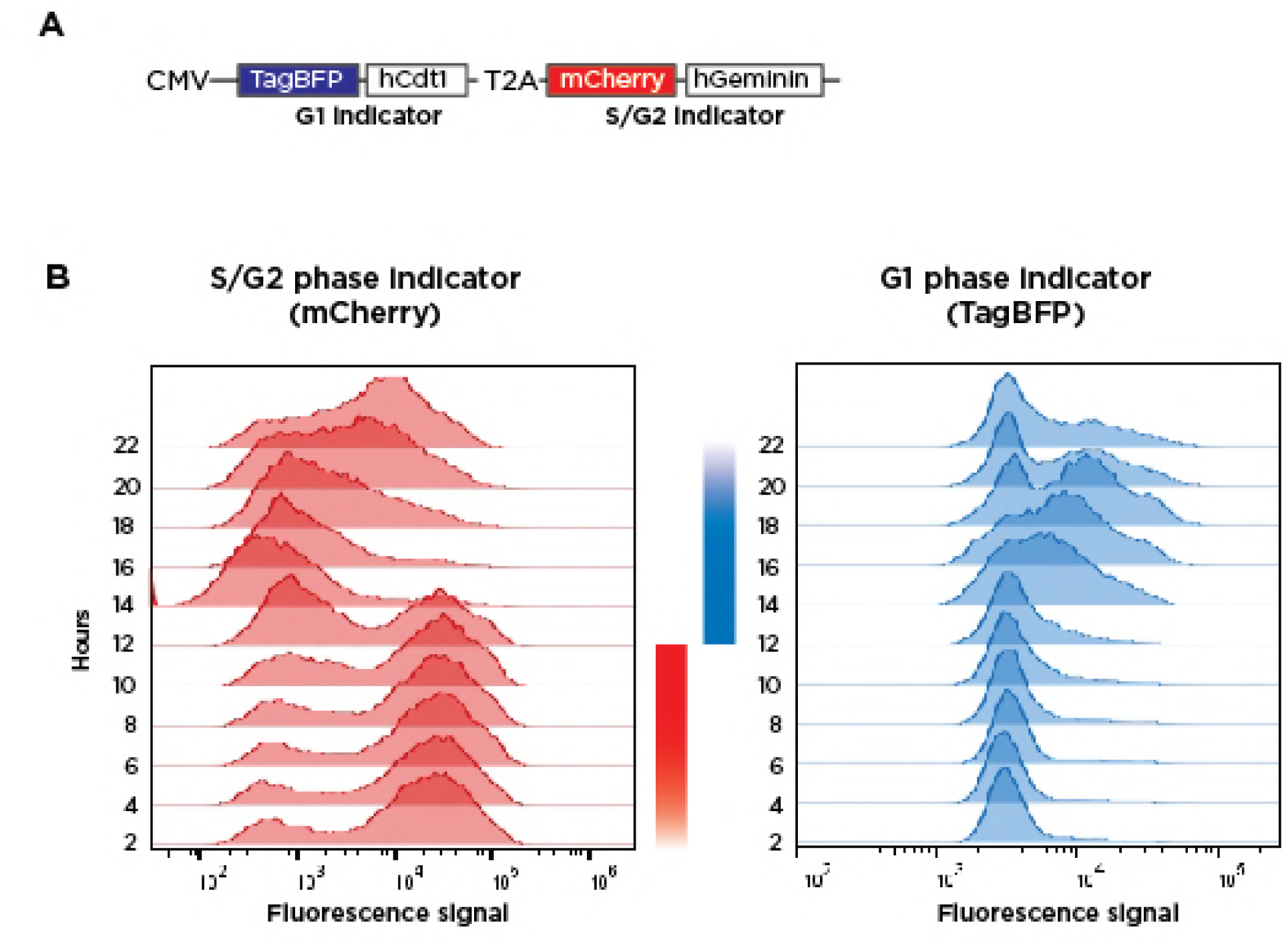
Determining the timing of the cell cycle in HEK 293T cells using flow cytometry. **(A)** The construct used for the fluorescence ubiquitination cell cycle indicator (Fucci (Sakaue-Sawano et al., 2008)). TagBFP-tagged hCdt1(30-120) and mCherry-tagged hGeminin (1-110) were expressed as G1 and S-G2 phase markers driven by a single UBC promoter using self-cleaving peptide T2A (*Thosea asigna* virus 2A) (Ryan et al., 1991). **(B)** Detection of the cell population in G1 (level of TagBFP) and S (level of mCherry) by flow cytometry. The cells were synchronized and released as described in **Figure 2**.

**Supplementary Figure S5:**
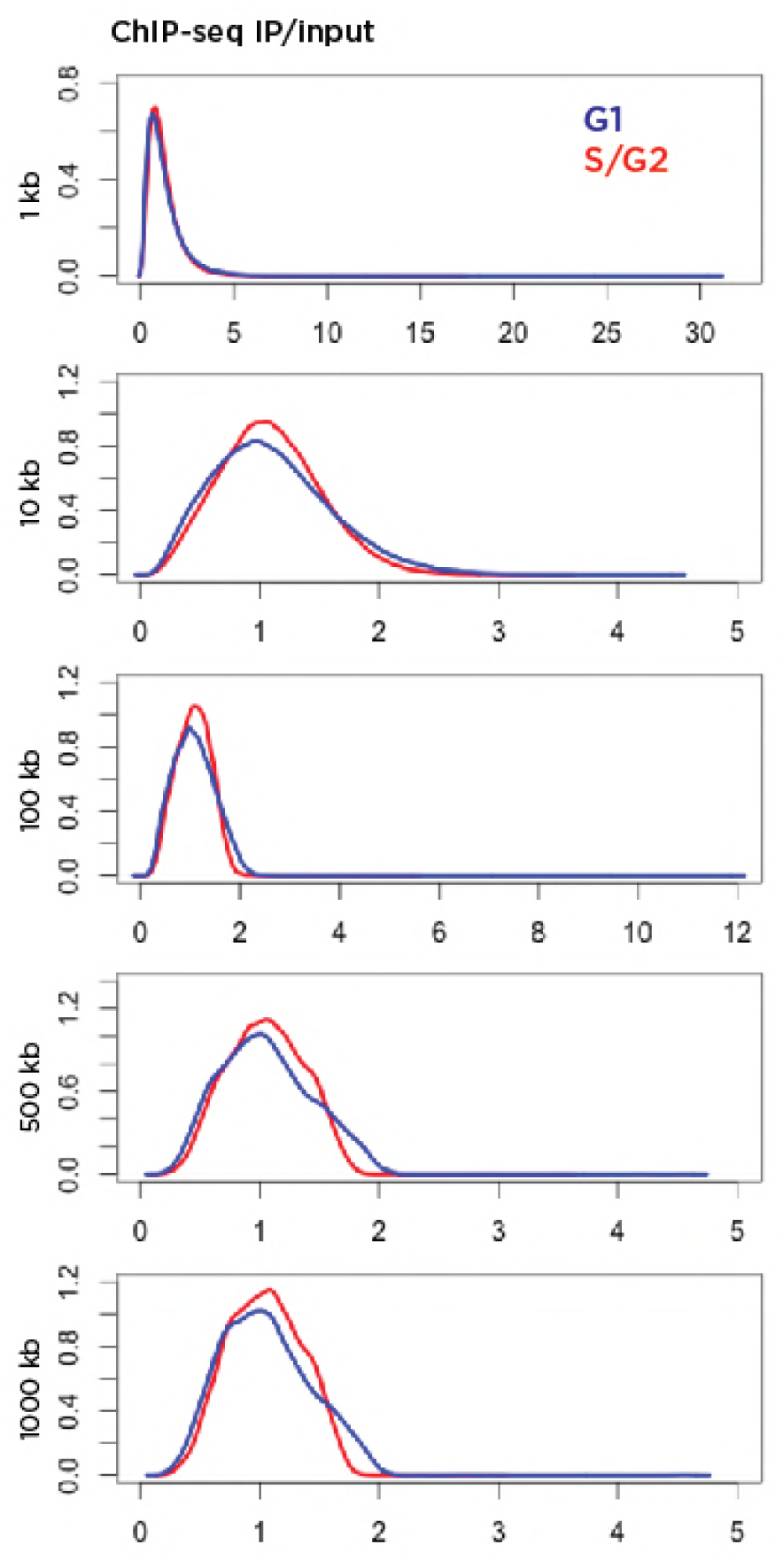
Analysis of the scale of enrichment for macroH2A ChIP-seq. Sequencing results of the immunoprecipitated (IP) SNAP-macroH2A (G1 or S/G2) and control input DNA were normalized to the same number of aligned reads per sample (∼127 million), with calculation of IP/input ratios for windows of 1 – 1,000 kb genome-wide. The top and bottom 1% of values were excluded and density plots generated as shown. No bimodality to suggest enrichment and specificity of targeting is apparent except in the windows of 500 – 1,000 kb.

**Supplementary Figure S6:**
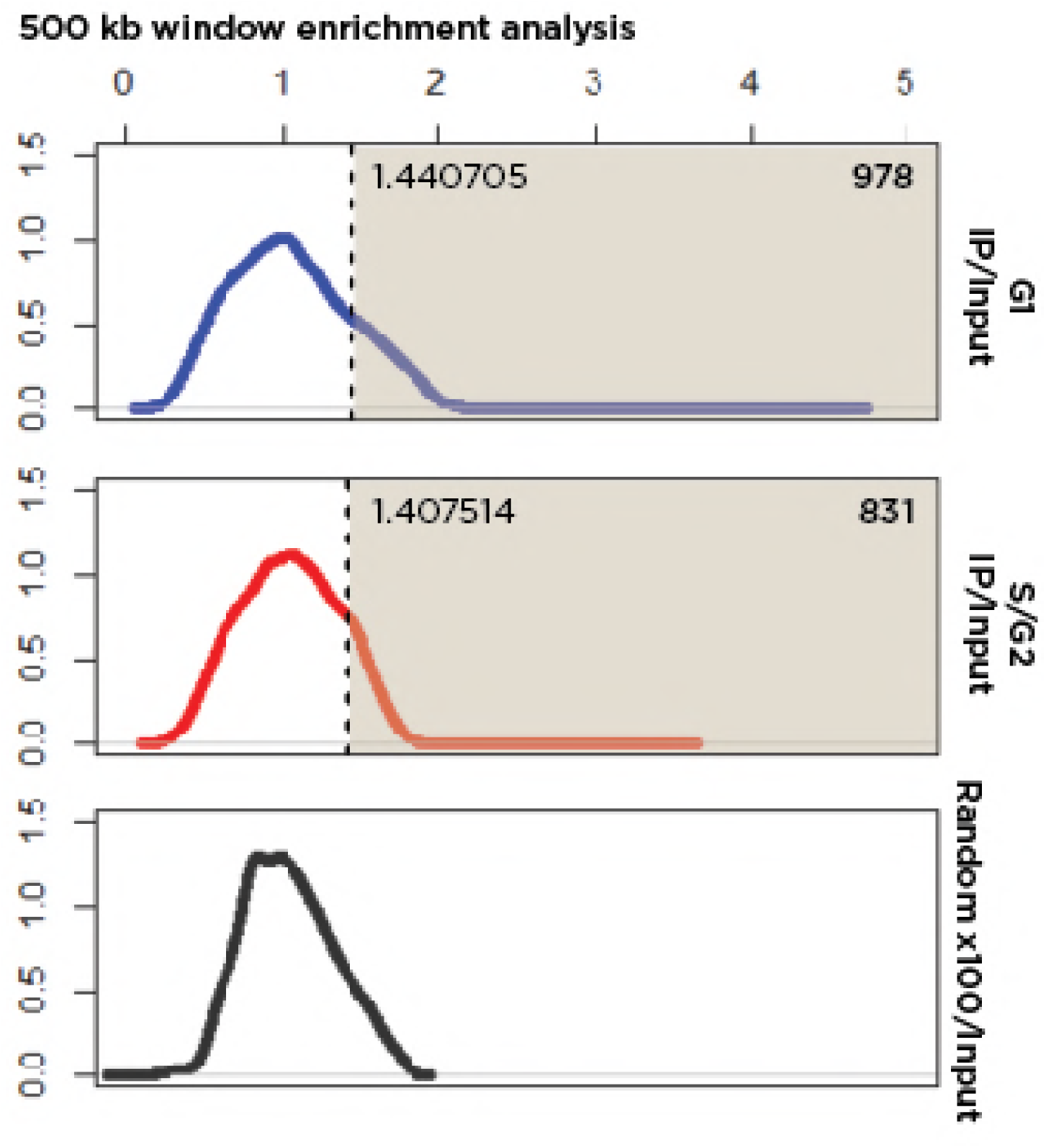
Identification of high-confidence loci of macroH2A enrichment in the genome. We used the 500 kb data as the smallest window revealing enrichment genome-wide (**Fig. S5**). The R package pastecs was used to identify the inflection point (pit, using turnpoints) where the second, enriched distribution begins. The cutoff values are shown for both the G1 and S/G2 distributions. The number of 500 kb windows identified to exceed these thresholds is shown in the top right of each plot. The bottom plot is the distribution of values following a random shuffling of window values 100 times, with no inflection point apparent for those data, indicating the specificity of the experimental results.

**Supplementary Figure S7:**
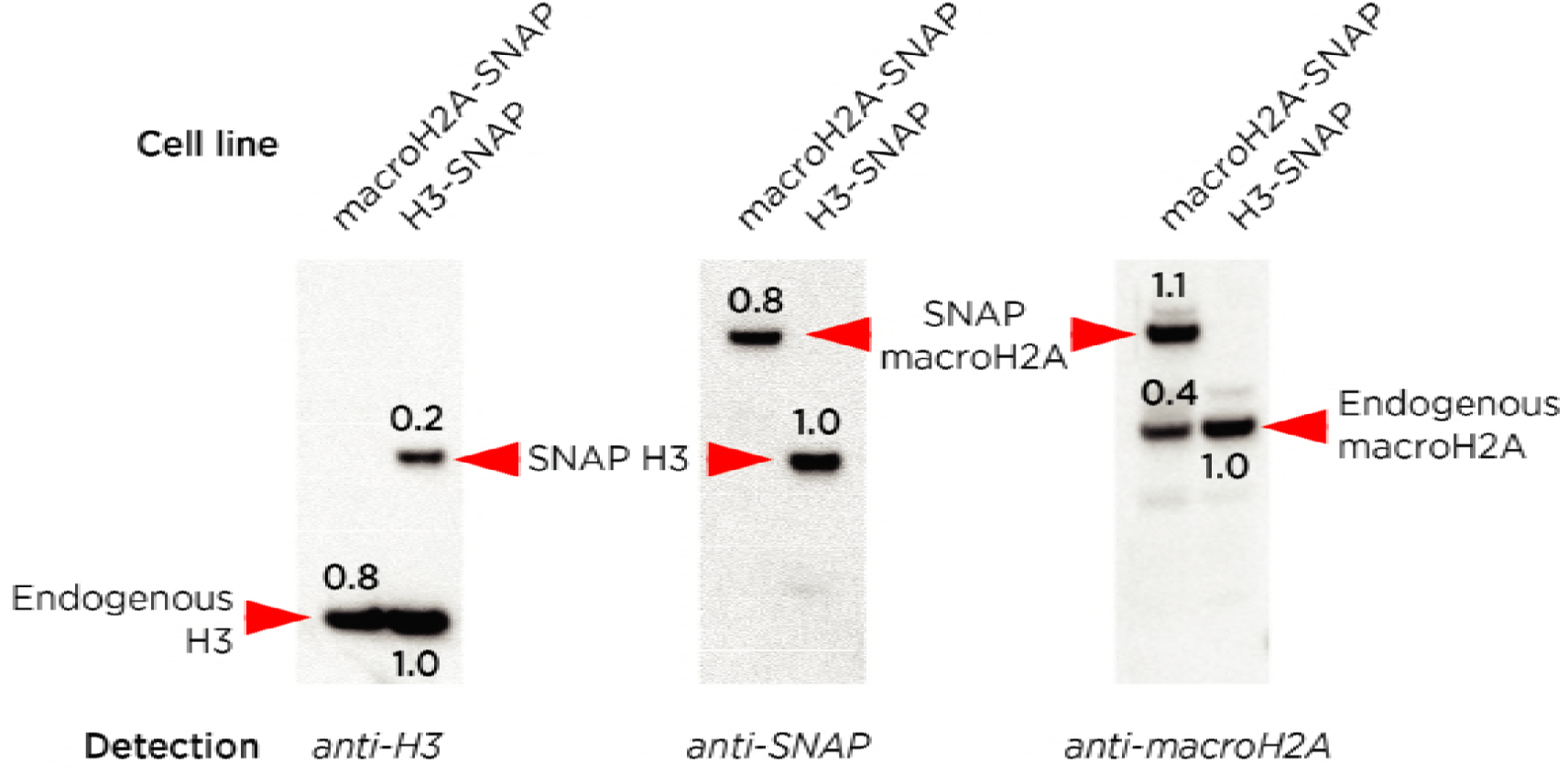
Quantification of proportion of nucleosomes with macroH2A. Mononucleosomes from HEK 293T cells stably expressing either SNAP-tagged macroH2A (left lanes) or histone H3 (right lanes) were loaded on a Western blot. Detection was performed using the antibodies shown under each image, and relative signal intensities measured by densitometry (relative values shown for each band). We assume that each mononucleosome contains a pair of histone H3 molecules, allowing the **left panel** to serve as a loading control. The right lane has a higher total signal intensity, (1.0 + 0.2 =) 1.2 / 0.8 = 1.5 fold that of the left lane, indicating the increased proportion of mononucleosomes loaded in the right lane. The **middle panel** reveals the differences in expression levels of the transgenes encoding the SNAP-tagged proteins, the macroH2A-SNAP expressed at (0.8 / (1.0 / 1.5) =) 1.2 fold the level of the H3-SNAP transgene. This allows us to test the proportion of mononucleosomes with endogeneous macroH2A in the H3-SNAP cell line (**right panel**). This endogeneous macroH2A occurs as a proportion of SNAP-macroH2A of (1.0 / 1.5) / 1.1, a ratio of 1:1.65. As macroH2A-SNAP is expressed at a ratio of 1.2: 1 of H3-SNAP, we calculate that endogeneous macroH2A is expressed at a ratio of 1: (1.65 / 1.20 =) 1.375 of H3-SNAP. From the **left panel**, we see that H3-SNAP is (0.2 / (0.2+1.0)=) 1/6 of all nucleosomes, allowing us to infer that endogeneous macroH2A is present in 1 in (1.375 x 6 =) 8.25, or 12.1%, of all nucleosomes.

**Supplementary Figure S8:**
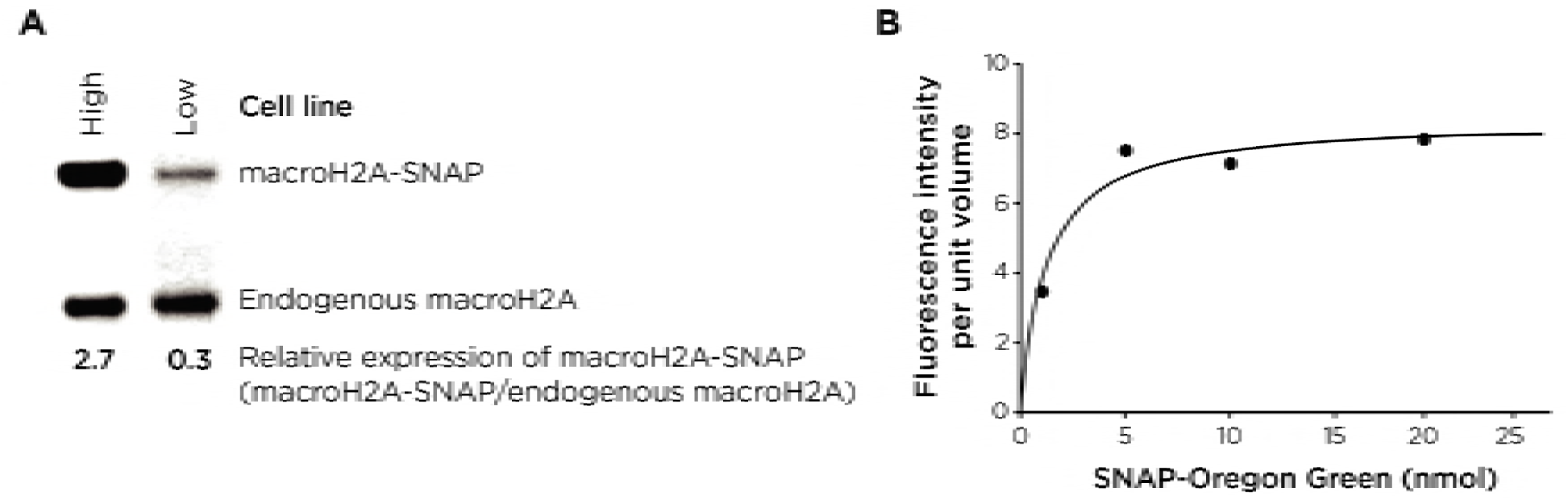
Conditions used for fluorescence correlation spectroscopy (FCS). The relative expression of SNAP-macroH2A relative to endogenous macroH2A in the HEK293T cell lines that were used in this study. We used the low-expressing line for all studies except for that involving FCS, for which we wanted to maximize the chances of incorporating two SNAP-macroH2A molecules per nucleosome. Titration of labeling SNAP-macroH2A with SNAP-Oregon Green for FCS analysis. The same amount of nuclei were used in labelling with 1, 5, 10, and 25 nmol of SNAP-Oregon Green and mononucleosomes labeled with SNAP-Oregon Green were detected by FCS. Values above 5 nmol of SNAP-Oregon Green saturate the fluorescence intensity values per molecule, indicating that the 5 nmol we used was occupying all available SNAP-tags.

**Supplementary Figure S9:**
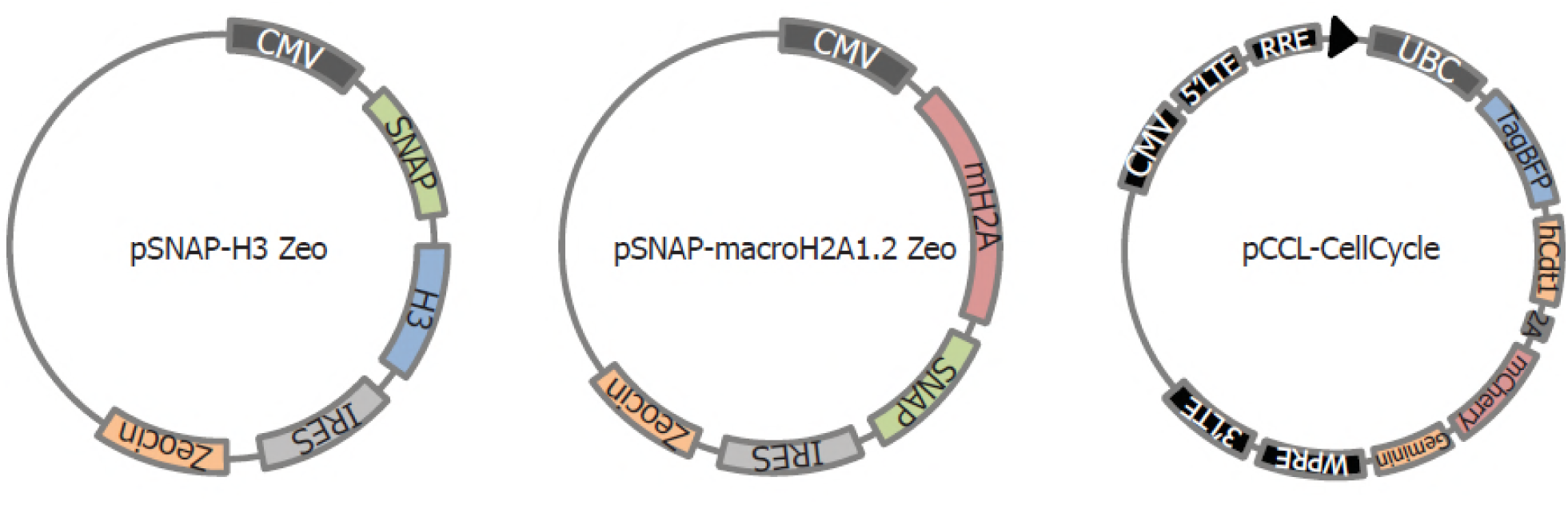
Plasmid constructs used. Due to the neomycin resistance of HEK 293T cells, which are immortalised with the large T antigen with neomycin selection, the neomycin drug selection marker in pSNAP_f_ (New England BioLabs) was replaced with Zeocin. cDNA of H3.1 or macroH2A1.2 (isolated from U2OS cells) was cloned into the N-terminal or C-terminal of SNAP-tag. The cell cycle indicator, containing G1 and S-G2 phase markers, was cloned into a lentiviral vector. Two protein, TagBFP-tagged hCdt1(30-120) and mCherry-tagged hGeminin (1-110) were expressed under single UBC promoter using self-cleaving peptide T2A (*Thoseaasigna* virus 2A).

## SUPPLEMENTARY TABLE

**Supplementary Table S1:**
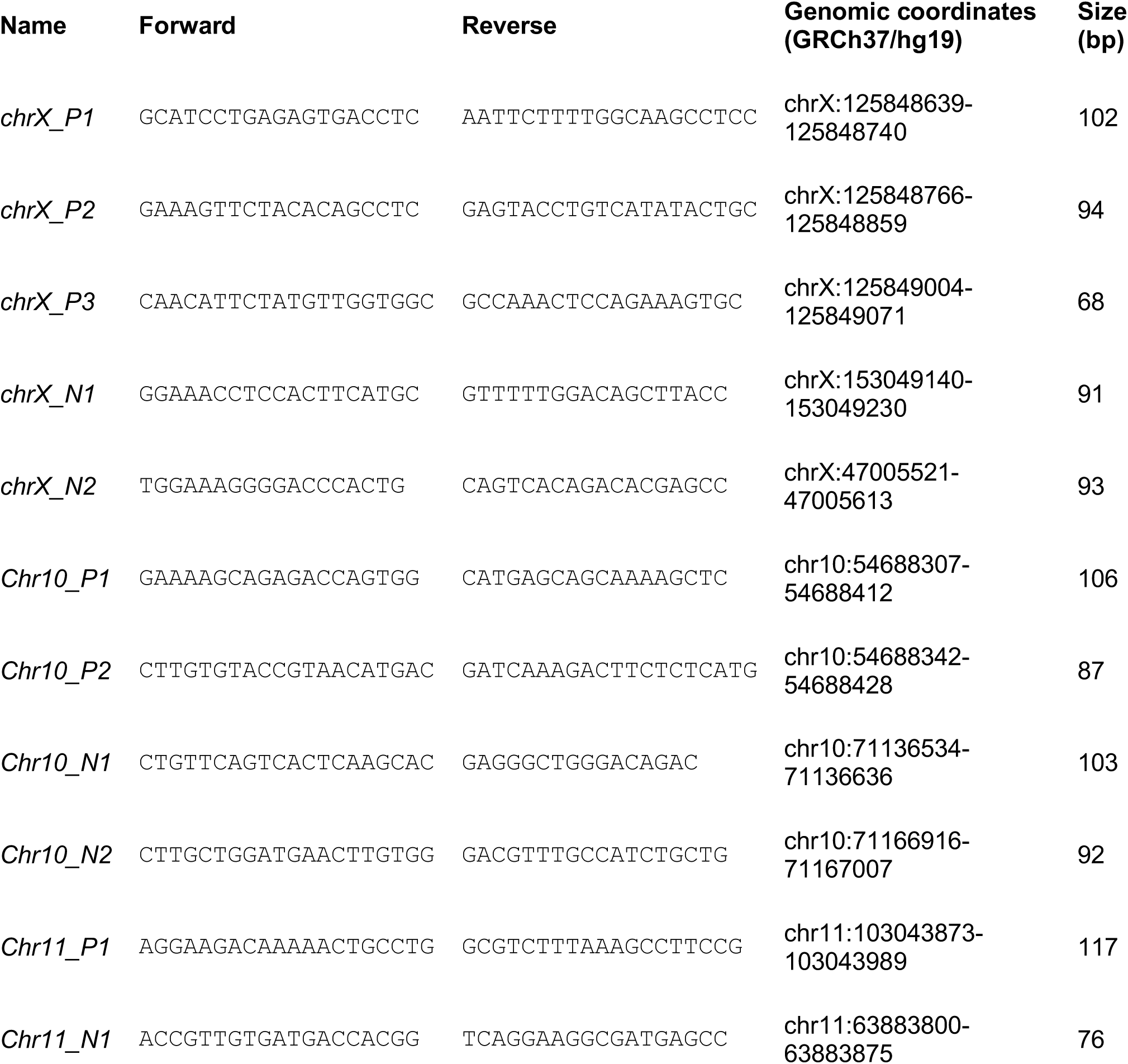
Primer list for ChIP-qPCR.

## KEY RESOURCES

| REAGENT or RESOURCE | SOURCE | IDENTIFIER |
| --- | --- | --- |
| **Antibodies** |  |  |
| Anti-Histone H3 Antibody | Abcam | cat#AB1791 |
| Histone macroH2A1 antibody | Active motif | cat#39593 |
| Histone H3K27me3 antibody | Active motif | cat#39155 |
| Anti-SNAP-tag® Antibody (Polyclonal) | New England BioLabs | cat#P9310S |
| Anti-phospho-Histone H3 (Ser10) Antibody, Mitosis Marker | Millipore | cat#06-570 |
| Monoclonal Anti-hnRNP-K/J antibody produced in mouse | Sigma | cat#R8903 |
| **Chemicals, Peptides, and Recombinant Proteins** |  |  |
| JF646 SNAP-tag Ligand | Grimm et al., 2015R.H. et al. Nat. Methods. 2015; 12: 244–250, 3, 250 | N/A |
| SNAP-Cell Oregon Green | New England BioLabs | cat#S9104S |
| SNAP-Cell Block | New England BioLabs | cat#S9106S |
| SNAP-Cell TMR-Star | New England BioLabs | cat#S9105S |
| SNAP-Biotin | New England BioLabs | cat#S9110S |
| Cell line Nucleofector Kit R | Lonza | cat#VCA-1001 |
| Thymidine | Sigma | cat#T1895 |
| RO-3306 | Santa cruz biotechnology | cat#sc-358700 |
| Nocodazole | Sigma | cat#M1404 |
| *Micrococcal nuclease* | Sigma | cat#N5386 |
| Prolong Diamond Antifade Mountant | Thermo Fisher Scientific | cat#P36970 |
| Prolong Diamond Antifade Mountant with Dapi | Thermo Fisher Scientific | cat#P36962 |
| Zeocin | Thermo Fisher Scientific | cat#R25001 |
| Pierce™ Protein A/G Magnetic Beads | Thermo Fisher Scientific | cat#88802 |
| Pierce™ Streptavidin Magnetic Beads | Thermo Fisher Scientific | cat#88817 |
| Power SYBR® Green PCR Master Mix | Thermo Fisher Scientific | cat#4367659 |
| **Critical Commercial Assays** |  |  |
| Click-iT® EdU Alexa Fluor® 647 Flow Cytometry Assay Kit | Thermo Fisher Scientific | cat#C10424 |
| SuperSignal West Pico Chemiluminescent Substrate | Thermo Fisher Scientific | cat#34080 |
| SuperSignal West Femto Chemiluminescent Substrate | Thermo Fisher Scientific | cat#34095 |
| Deposited Data |  |  |
| ChIP-Seq | This paper | NCBI SRA: |
| Scripts for ChIP-seq analysis | This paper | https://github.com/hnst/Sato-et-al |
| Macro for analysis of imaging acquisition | This paper | https://github.com/hnst/Sato-et-al |
| Experimental Models: Cell Lines |  |  |
| Human: HEK293T | ATCC | cat#CRL-3216 |
| **Recombinant DNA** |  |  |
| pSNAPf Vector | New England BioLabs | cat#N9183S |
| pSNAPf-Histone H3.1 zeo | This paper | N/A |
| pSNAPf-macroH2A1.2 zeo | This paper | N/A |
| phage-CellCycle | This paper | N/A |
| pCS2+ macroH2A1.2-mCherry-FLAG | This paper | N/A |
| Sequence-Based Reagents |  |  |
| PCR Primers: ChIP-qPCR, see **Supplementary Table S1** | This paper (IDT) | N/A |
| **Software and Algorithms** |  |  |
| FlowJo | FlowJo, LLC | https://www.flowjo.com/ |
| ImageJ | NIH | https://imagej.nih.gov/ij/ |
| GraphPad Prism 7 | GraphPad Software Inc | http://www.graphpad.com/scientific-software/prism/ |
| Integrated Genomics Viewer (IGV) | Robinson et al., 2011; Thorvaldsdottir et al., 2013 | https://www.broadinstitute.org/igv/ |
| BWA | Li H. and Durbin R. (2009) | http://bio-bwa.sourceforge.net/ |
| SAMtools | Li et al., 2009 | http://samtools.sourceforge.net |
| Picard-tools | MIT | http://picard.sourceforge.net |
| Bedtools2 | Quinlan 2014 | http://code.google.com/p/bedtools/ |
| R | R Development Core Team | https://www.r-project.org/ |

## CONTACT FOR REAGENT AND RESOURCE SHARING

For the reagents generated in this study please contact John M. Greally

## METHODS

### Cell Lines and Tissue Culture

HEK293T cells were grown at 37°C and 5% CO_2_ in DMEM containing 4.5 g/l of glucose, 10% FBS and 1% penicillin-streptomycin. HEK293T cells stably expressing SNAP-H3 or macroH2A1.2 were selected with Zeocin (100 nM) treatment for 3 weeks after transfection of linearized vector. HEK293T stably expressing cell cycle marker was established by lentivirus packaging system as previously described (Naldini et al., 1996). The cells expressing each gene were selected by fluorescence-activated cell sorting (FACS).

### Plasmid Construction

In order to generate the plasmids expressing SNAP-H3 and SNAP-macroH2A, the pSNAP_f_ vector produced from New England BioLabs was used. Due to the drug incompatibility in HEK293T cells, we replaced the existing Neomycin selection gene with Zeocin. After creating the pSNAP_f_-Zeocin vector, the cDNA of H3.1 isolated from a human cDNA library or the cDNA for macroH2A1.2 from pCS2+ macroH2A1.2-GFP-HA (Addgene #30515) were cloned into the pSNAP_f_-Zeocin vector. The Fucci cell cycle indicator (Sakaue-Sawano et al., 2008), which expresses TagBFP-tagged hCdt1 (30-120) and mCherry-tagged hGeminin (1-110) with a T2A self-cleaving peptide T2A (*Thosea asigna* virus 2A) (Ryan et al., 1991) was cloned into a lentiviral vector. The maps of all constructs are shown in **Supplementary Figure S9**. For the construct expressing mCherry-tagged macroH2A1.2, results shown in **Supplementary Figure S1**, GFP was replaced by mCherry in pCS2+ macroH2A1.2-GFP-HA (Addgene #30515).

### Synchronization of cell cycle, Histone labeling using SNAP tag and EdU incorporation

Cells were synchronized at the G1/S transition using the double thymidine block method, using a treatment of 2 mM thymidine for 16 hours, releasing from thymidine for 9 hours and treating again with 2 mM thymidine again for 17 hours. SNAP-tagged histones were labeled with cell permeable SNAP-substrates, SNAP-Cell Oregon Green (1 µM), SNAP-Cell TMR (1 µM) or blocked with non-fluorescent SNAP-substrates, SNAP-cell Block (10 µM), for 30 minutes at 37°C in 5% CO_2_. Proliferating cells were detected using the Click-iT^®^ EdU Cell Proliferation Assay (Life Technologies). To collect mitotic cells, a mitotic shake off was performed 12 hours after treatment with 20 nM Nocodazole, following treatment with 2 mM thymidine for 24 hours.

### Fixed cell image acquisition

For imaging of fixed cells, cells were grown on poly-L lysine-coated coverslips and fixed in 4% paraformaldehyde for 15 minutes at room temperature, then permeabilized with 0.2% Tween in PBS for 15 minutes at room temperature. For the immunofluorescence experiment in **Supplementary Figure S1**, the cells were treated in PBS containing 3% BSA for 1 hour, then incubated with primary antibody (1/1000 dilution) overnight. After washing with PBS three times, the cells were incubated with secondary antibody (1/10,000 dilution) for 30 minutes. The cells were mounted with Prolong Diamond Antifade Mountant with DAPI. Images were acquired with an Olympus BX61 widefield, epifluorescent microscope using a 60X 1.4 PlanApo objective. Filter sets were used for DAPI (Semrock model DAPI-5060C-Zero), Cy3 (Chroma model 41007), FITC (Semrock model FITC-5050A-Zero), and Cy5 (Semrock model Cy5-4040C-Zero), with an EXFO X-Cite Series 120 PC metal halide light source, Photometrics Cool SNAP HQ CCD camera, Olympus Type-F immersion oil (nd 1.516) and Molecular Devices Metamorph acquisition software. Cells were optically sectioned using a 0.5 μm Z step, spanning a 3.0 μm Z depth in total. Exposure times of 20 to 200 msec were typically used to acquire each plane in the Cy3, Cy5 and FITC channels, and ∼12 ms were used to acquire each plane in the DAPI channel.

### Live cell imaging acquisition

For live cell imaging, we replaced the growth medium with FluoroBrite(tm) DMEM Media containing 10% FBS, 1% penicillin/streptomycin and GlutaMax before imaging. Wide-field images of mitotic cells were taken on an IX-81 stand (Olympus). The microscope was equipped as described previously (Wu et al., 2016). The cells were kept at 37°C with a stage top incubator (INUBH-ZILCS-F1, Tokai Hit, Japan) in 5% CO_2_. Cells were optically sectioned using a 500 nm Z step, spanning a 5.0 μm Z depth in total. 50 msec exposure times were used to acquire each plane in all channels.

### Cell cycle specific native ChIP using SNAP-Biotin

A total of 1-2 x10^7^ cells were suspended in 1 ml extraction buffer (10 mM HEPES pH 7.5, 10 mM KCl, 1.5 mM MgCl_2_, 0.34 M Sucrose, 10% Glycerol, 0.2% NP40, Protease inhibitor) after treatment with the SNAP-block approach, as described for **Supplementary Figure 3A**, then lysing the cells for 10 minutes on ice. Nuclei were pelleted (6,500 rpm, *5 minutes, 4°*C) and the supernatant discarded. Isolated nuclei were washed with 1 ml of extraction buffer without NP40 and suspended in 0.5 ml digestion buffer (50 mM Tris pH 7.5, 1 mM CaCl_2_, 1 mM DTT, 1 mM PMSF, Protease inhibitor). The DNA concentration was detected using 2 M NaCl adjusting the concentration of the chromatin fraction to 1µg/ml. Newly-incorporated macroH2A in 1 ml of nuclei (1 µg/ml) were labeled with SNAP-Biotin (1 µM) at 4 °C for 30 minutes. Ten microliters of 50 mU/µl *Micrococcal nuclease* (Sigma) were added and incubated at 37 °C for 10 minutes. The reaction was stopped by supplementing with 100 µl of 0.1 M EGTA. Digested chromatin was centrifuged at 8,000 g at 4 °C for 5 minutes. Supernatant was labelled as S1 and stored at 4 °C, and the pellet was suspended in lysis buffer (Tris pH 7.4, 0.2 M EDTA, Protease inhibitor) and dialyzed against 2 L of dialysis buffer (Tris pH 7.4, 0.2 mM EDTA) overnight. The chromatin fraction was centrifuged at 500xg for 10 minutes at 4 °C, and the supernatant was combined with the S1 sample, keeping 100 µl of this fraction as our Input sample. Fifty microliters of streptavidin magnet beads pre-blocked with 1 mg/ml BSA and 0.3 mg/ml Salmon Sperm DNA were added to 150 µg of the chromatin fraction and rotated for 2 hours at 4 °C. Beads were washed five times and DNA isolation was performed followed by ChIP-seq library preparation as we have described previously (Ramos et al., 2015), sequencing using the Illumina HiSeq 2500 generating 100 bp paired-end reads, and validating results using ChIP-qPCR using the Power SYBR^®^ Green PCR Master Mix (Thermo Fisher Scientific). The levels of fold-enrichment were determined from *C*T values normalized by negative control of immunoprecipitated with non-specific rabbit IgG. The list of primers is shown in **Supplementary Table S1**.

### Mononucleosome isolation and immunoprecipitation

Chromatin isolation and MNase digestion were performed as described in the previous section. The salt concentration in the nucleosome fraction was adjusted to be 0.65 M NaCl, applied on the top of a 5-28% sucrose gradient (20 mM Tris pH7.5, 150 mM NaCl, 0.2 mM PMSF) and ultracentrifuged for 15 hours, 36,000 rpm at 4 °C using a Beckman SW41 rotor. Nucleosomes were fractionated by taking samples sequentially from the top. DNA was purified from an aliquot from each fraction and mono-nucleosome fractions were identified by the size of DNA fragments. The purified mononucleosome fraction was incubated with 5 µl of anti-SNAP antibody (New England BioLabs) or rabbit IgG for 2 hours at 4 °C. Fifty microliters of protein A/G magnet beads (Thermo Fisher Scientific, #26162) pre-blocked with 1 mg/ml BSA were added and rotated for 1 hour at 4 °C. The beads were washed three times and proteins were eluted by SDS loading buffer.

### Fluorescence fluctuation microscope (FFS)

For FFS analysis, 150 µl of purified nuclei (1 mg/ml) were labeled with 5 nmol of SNAP-OG for 2 hours at RT followed by MNase digestion and ultracentrifugation as described above. The mononucleosome fraction was measured on 5% BSA coated 8-well chambered coverglass (Thermo Fisher Scientific, 155411) using a home-built two-photon fluorescence fluctuation microscope described previously (Wu et al., 2015).

## DATA ANALYSIS

Quantifications and statistical analyses in imaging acquisitions and in Western blotting were performed using ImageJ and GraphPad Prism 7. For the analyses of **Supplementary Figures 1A** and **1D**, two-tailed unpaired t-tests were performed in GraphPad Prism 7. Quantifications and statistical analyses in FACS analyses were performed in FlowJo.

## DATA AND SOFTWARE AVAILABILITY

The ChIP-seq data reported in this paper are available from the NCBI Sequence Read Archive (SRA), accession number SRP092259.

https://trace.ncbi.nlm.nih.gov/Traces/sra/sra.cgi?study=SRP092259

Reviewer link to metadata:

ftp://ftp-trace.ncbi.nlm.nih.gov/sra/review/SRP092259_20180504_172013_cb5ae17636e975f9bf71ddf5bc542075

Custom code used in this project is available at GitHub https://github.com/hnst/Sato-et-al

